# Elucidating the effect of a rationally designed nanostructured form-switching ASO (NaFASO) for targeting long non-coding RNA to alleviate Japanese encephalitis virus infection

**DOI:** 10.64898/2026.05.30.728934

**Authors:** Chandrika Sharma, Suryansh Sengar, Dharani Sen, Vivek Sharma, Souradyuti Ghosh

**Author notes:** Joint corresponding author. Equal contribution as co-first authors (Chandrika Sharma, Suryansh Sengar). Will handle communication (,) (Souradyuti Ghosh).

## Abstract

RNA therapeutic modalities such as antisense oligonucleotides (ASOs) have emerged as promising tools to target previously “undruggable” targets. Despite their great promise as precision therapeutic agents, their clinical adoption remains limited due to production costs, sequence-length restrictions, limited structural heterogeneity, and the generation of environmentally hazardous waste during synthesis. Biocatalytic synthesis strategies provide a sustainable alternative; however, their reliance on specialized enzymes and precursors often limits sequence diversity and scalability. To address these limitations, we report the design and biocatalytic synthesis of a novel circular ASO: Nanostructured Form-switching Antisense Oligonucleotide (NaFASO) for targeting Japanese Encephalitis Virus (JEV) infection-associated host long non-coding RNA (lncRNA) *JINR1* (JEV-induced non-coding RNA1) in SH-SY5Y cells. The novel modular architecture in NaFASO has been designed to have a metastable stem that separates the functional antisense domain from the splint-padlock circularizing region, ensuring both structural integrity and efficient target engagement. The serum- and nuclease-stable NaFASOs achieved knockdown of the lncRNA *JINR1* during JEV infection, resulting in a reduction in JEV replication and neuronal cell death. NaFASO-mediated *JINR1* depletion also resulted in downregulation of the JEV replication-associated gene GRP78. Together, these findings establish NaFASO as a first-of-its-kind structure-switching circular ASO platform for combating JEV infection, combining stability, efficacy, and environmental sustainability. Beyond the JEV, the generalizability of this design suggests broad applicability for targeting diverse RNA species implicated in genetic disorders, viral infections, and cancer, thus highlighting a promising paradigm for developing next-generation transformational nucleic acid therapeutics.

## Introduction

Oligonucleotide-based therapeutics have emerged as one of the most versatile and rapidly advancing modalities in molecular medicine, offering a precise and programmable approach to modulate gene expression by regulating transcript amounts^1^. These agents, which include ASOs, small interfering RNAs (siRNAs), and aptamers, function by hybridizing to complementary RNA targets to induce degradation, alter splicing, or block translation^2–4^. Unlike small-molecule drugs or protein biologics, oligonucleotides can be rationally designed solely from sequence information, enabling the treatment of diseases previously considered “undruggable” ^5^. Several ASO-based drugs—such as *Nusinersen* for spinal muscular atrophy and *Eteplirsen* for Duchenne muscular dystrophy—have validated the clinical potential of this technology^6,7^.

Backbone-modified ASOs have been extensively developed to address instability and degradation in biological systems. Locked nucleic acids (LNAs), phosphorodiamidate morpholino oligomers (PMOs), and phosphorothioate derivatives are among the most widely used backbones that enhance nuclease resistance and binding affinity in vivo^8–10^. However, these chemical modifications are accompanied by significant drawbacks. Their synthesis requires multi-step solid-phase chemistry that is chemically intensive, generates organic waste, and limits scalability^11,12^. Furthermore, the incorporation of non-natural backbones may alter pharmacokinetic and pharmacodynamic profiles, elicit unintended immune responses, and complicate regulatory evaluation^10,13^. In addition, unmodified nucleic acids remain susceptible to exonuclease-mediated degradation, especially at their 5′- and 3′-termini, resulting in short half-lives within cells^13–15^. Consequently, there is growing interest in alternative design strategies that could provide intrinsic nuclease resistance while maintaining biocompatibility and minimizing the reliance on chemically modified backbones.

Circularization of oligonucleotides is a promising strategy to circumvent nuclease degradation without requiring chemical modification^16^. By covalently linking the 5′- and 3′-termini to form a closed circular structure, these molecules effectively eliminate free ends, rendering them highly resistant to exonuclease attack^17–19^. Circular oligonucleotides have thus demonstrated remarkable stability, improved hybridization kinetics, and enhanced persistence within cellular environments. Several chemical and enzymatic approaches—such as splint-padlock ligation, click chemistry, and T4 DNA ligase-based circularization—have been reported for their synthesis^20–22^. Despite these successes, most existing chemical circularization methods involve complex multi-step reactions, use costly reagents, and suffer from low yields and limited scalability ^23,24^. Moreover, integrating functional antisense domains into circular frameworks without compromising folding or hybridization efficiency remains technically challenging ^25^. Thus, there is a strong need for a bio-catalytic, environmentally sustainable, and modular synthesis route that can produce circular oligonucleotides with tuneable sequence designs and high functional efficacy ^26^.

Neurotropic viral infections pose a particularly critical and unmet clinical challenge. JEV is a mosquito-borne neurotropic virus that belongs to the *Orthoflavivirus* genus^27^. It continues to cause more than 68,000 clinical cases annually across South and South-east Asia, with a fatality rate of 20–30% and long-term neurological sequelae in survivors^28–32^. Despite the availability of vaccines, outbreaks persist due to incomplete immunization coverage ^33–37^. Moreover, there is no approved small-molecule therapy against JEV infection. Host lncRNAs play critical role in regulating JEV replication and inflammation ^38,39^. LncRNA-*JINR1* (also known as LINC01518) regulates JEV replication and associated cell death^40^. Knockdown of *JINR1* using chemically modified ASOs (LNA ASOs) markedly reduces viral replication, cell death and expression of GRP78, a gene involved in JEV entry and replication in host cells^39^. However, the cost, complexity, and limited scalability of such backbone-modified oligonucleotides restrict their translational potential for widespread antiviral applications.

In this context, the present work introduces Nucleic acid Functionalized Antisense Oligonucleotide (NaFASO), a novel circular form-switching DNA nanostructural construct synthesized through an enzymatic splint–padlock ligation. This bio-catalytic approach eliminates the need for synthetic backbone modifications while achieving functional efficiency alongside maintaining structural stability in serum and nuclease environments. Designed with distinct stem, antisense, and splint-binding regions, NaFASO ensures the precise hybridization of the functional antisense domain to its RNA target while maintaining the integrity of its secondary structure. Using the JEV-infected SH-SY5Y cell line as a model system, we demonstrate a proof-of-concept NaFASO-mediated knockdown of lncRNA *JINR1* during JEV infection, which reduces JEV replication and associated cell death. The system offers an effective combination of stability, selectivity, and ease of biosynthetic preparation, establishing NaFASO as a robust and scalable platform for antisense therapeutics.

## Experimental

### General

All oligonucleotides used in this study (sequences listed in Table S1) were purchased from Eurofins Genomics and supplied following high-purity salt-free (HPSF) purification. The oligonucleotides were used without further purification. T4 polynucleotide kinase (M0201S), T4 DNA ligase (M0202SVIAL), Exonuclease I (M0293S), Exonuclease III (M0206SVIAL), bovine serum albumin (BSA, P8108S), adenosine 5′-triphosphate (ATP, P0756L), T4 DNA ligase buffer (B0202SVIAL), and NEB buffer 1 (B7001SVIAL) were obtained from New England Biolabs (USA). Tris base (71033), spermidine (17000), and dithiothreitol (DTT, 17315) were purchased from Sisco Research Laboratories Pvt. Ltd. (SRL), India. Phosphodiesterase I from Crotalus adamanteus venom (P3243-1VL) was procured from Sigma-Aldrich. Fetal bovine serum (FBS), Dulbecco’s Modified Eagle Medium (DMEM), DMEM/F12 medium, Oxoid phosphate-buffered saline (BR0014G), and paraformaldehyde (Q23996) were purchased from Thermo Fisher Scientific. Penicillin–streptomycin solution (9802000) was obtained from Invitrogen. Agarose MP (11388983001) and crystal violet (V5265) were acquired from Sigma-Aldrich. All aqueous solutions were prepared using ultrapure water from a Milli-Q Type II water purification system (Millipore). Experimental procedures were carried out using a Prima-96 thermal cycler (HiMedia), a SpectraMax iD5 microplate reader (Molecular Devices), and a CFX Opus 96 Real-Time PCR system (Bio Rad).

### Phosphorylation

Phosphorylation was carried out in 100 μL reaction volume in the presence of T4 PNK, buffer (50 mM Tris-HCl pH 7.5, 10 mM MgCl_2_, 1 mM ATP, 10 mM DTT), and water to make up the volume. The solution was incubated at 37°C for 3 h, followed by inactivation by heating at 75°C for 20 min. Afterwards, the splint was mixed with the 5’-phosphorylated padlock, followed by annealing.

### Ligation

Circularization of the annealed 5’-phosphorylated DNA was carried out in the presence of T4 DNA ligase, ATP, and T4 DNA ligase (50 mM Tris-HCl pH 7.5, 10 mM MgCl_2_, 1 mM ATP, 10 mM DTT) buffer in 100 μL reaction volume. The ligation mixture was then incubated at room temperature (25 °C) for 4 h, followed by enzyme inactivation at 75°C for 20 min.

### Exonuclease digestion

The exonuclease digestion was performed in a 100 µL reaction volume, as described below. For exonuclease digestion, the reaction was carried out with both the exonuclease enzymes in buffer (10 mM Bis-Tris-Propane-HCl, 10 mM MgCl_2_, 1 mM DTT, pH 7). The reagents were added while keeping the vials on ice. The solutions were incubated at 37 °C for 4 h, followed by enzyme inactivation at 85 °C for 20 min. The sample concentration was quantified by densitometric analysis of sample bands in 12% Urea PAGE to confirm the final product’s concentration.

### Snake venom phosphodiesterase (SVPD) Assay

250 ng of linear DNA or NaFASO were incubated with SVPD (final 2 mU/μL) in DMEM 10% FBS media at 37 °C for time points of 0, 5, 15, 30, 60, and 90 minutes. The reaction was terminated by heating the sample at 80 °C for 5 min. Sample stability was determined by densitometric analysis of sample bands, along with an untreated 250 ng control, in 12% Urea PAGE.

### Cell culture

SH-SY5Y cells were cultured in DMEM/F12 medium supplemented with 10% heat-inactivated fetal bovine serum (FBS), 2 mM L-glutamine, and penicillin–streptomycin (Gibco) under standard conditions (37 °C, 5% CO_2_). PS and Vero cells were maintained in Dulbecco’s modified Eagle medium (DMEM) supplemented with 10% heat-inactivated fetal bovine serum (FBS), 2 mM glutamine, and penicillin/streptomycin (Gibco) at 37°C and 5% CO_2_.

### Virus propagation and Plaque Assay

The GP78 strain of JEV was propagated in Vero cells as described previously^40,41^. Briefly, Vero cell monolayers were inoculated with JEV at 0.1 MOI and incubated at 37°C and 5% CO_2_ for 72 hpi (hours post-infection) until the infected monolayers showed cytopathic effects. Subsequently, the culture supernatant was harvested by centrifugation at 2,000 rpm for 20 min at 4°C, then filtered through 0.8/0.2 µm syringe filters (Acrodisc Syringe Filters, #4658) and aliquoted to generate a virus stock. JEV titer was determined by plaque assay in PS cells. All virus stocks were aliquoted and stored at −80°C.

The plaque assay was performed using PS cells. For plaque assays, 2 × 10^5^ PS cells were seeded into 6-well culture plates. After 24 h, the monolayers were infected with serially diluted cell culture-grown JEV (GP78) for 2 h. After adsorption, PS cells were washed twice with PBS to remove unbound virus, then overlaid with a 1:1 mixture of 2% low-melting agarose and growth medium. Cultures were incubated for 4 days at 37 °C in 5% CO_2_, after which they were fixed in 4% formaldehyde for 1 h at 37 °C. Plaques were visualized by staining with 0.5% crystal violet solution for 5 min, followed by three washes with RO water. Viral titers were determined as plaque-forming units (PFU) per millilitre according to the formula: PFU = (N × DF)/V, where *N* is the number of plaques, *DF* the dilution factor, and *V* the inoculum volume.

### NaFASO Transfection

SH-SY5Y cells were reverse-transfected with NaFASOs using Lipofectamine RNAiMAX (Invitrogen, #13778-075) according to the manufacturer’s protocol. Briefly, 0.8×10^6^ cells were reverse-transfected with 60 pmol NaFASO, followed by virus infection 18 hours post-transfection.

### Viral infection

One day post-seeding, SH-SY5Y cells were serum-starved for 2 h and infected with JEV at an MOI of 5 in incomplete DMEM-F12 medium at 37°C for 2 h. After virus adsorption, the inoculum was removed, and the cells were washed with 1X PBS. After removing the virus-containing medium, the cells were grown in reduced-serum (5%) growth media for 48h, followed by RNA extraction.

### Cell Proliferation Assay

Cell proliferation was evaluated using the WST-1 reagent (Roche, #05015944001) according to the manufacturer’s instructions. Briefly, SH-SY5Y cells were seeded into 96-well plates and reverse-transfected with NaFASO Scrambled JINR1-S (control), NaFASO JINR1-1, and NaFASO JINR1-2 at a final concentration of 60 pmol. Cell proliferation was measured by recording absorbance at 450 nm.

### Caspase Assay

Caspase-3/7 activity was measured using a luminometric assay kit (Promega, #G8090) according to the manufacturer’s protocol. SH-SY5Y cells were transfected with NaFASO scrambled JINR1-S (control), NaFASO-JINR1-1, and NaFASO JINR1-2, and subsequently, luminescence was recorded at the indicated time point to measure the relative caspase 3/7 activity.

### RNA isolation and real-time PCR

Total RNA was isolated using the Direct-zol RNA Miniprep Kit (Zymo, #R2050). Subsequently, 1 µg of RNA was reverse transcribed into cDNA with the PrimeScript First-Strand cDNA Synthesis Kit (Takara, #6110A). Quantitative real-time PCR (qRT-PCR) was carried out using the SYBR Green PCR Kit (Takara, #RR820A) on a CFX Opus 96 Real-Time PCR system (Bio Rad). Each reaction was performed in triplicate, with GAPDH serving as the internal reference gene. Relative expression levels were determined using the 2^-ΔΔCt^ method.

## Results

### Design consideration of NaFASO

We synthesized two distinct NaFASO constructs targeting lncRNA *JINR1*, namely NaFASO JINR1-1 and NaFASO JINR1-2 (Supplementary Table S1), along with a scrambled control, JINRI-S. The antisense sequences for NaFASO JINR1-1 and NaFASO JINR1-2 were derived from a study by Tripathi et al, which involved LNA-based knockdown of *JINR1*^40^. In our construct, NaFASO was engineered to integrate three functional domains (Fig. 1A – D and Fig. S1): (i) ASO region for sequence-specific target recognition, (ii) a flexible 5 nt stem region to bring the termini into proximity, and (iii) a splint-binding region that enables efficient ligation-mediated circularization. Precise sequence design was essential, as suboptimal base composition or secondary structure could compromise both ligation efficiency and the specificity of the target interaction (Supplementary Table S1). The oxView simulations further corroborated that the designed NaFASO JINR1-1 (Fig. 1C) and NaFASO JINR1-2 (Fig. 1D) constructs adopt the intended structural configuration upon circularization^42^.

**Fig. 1.**
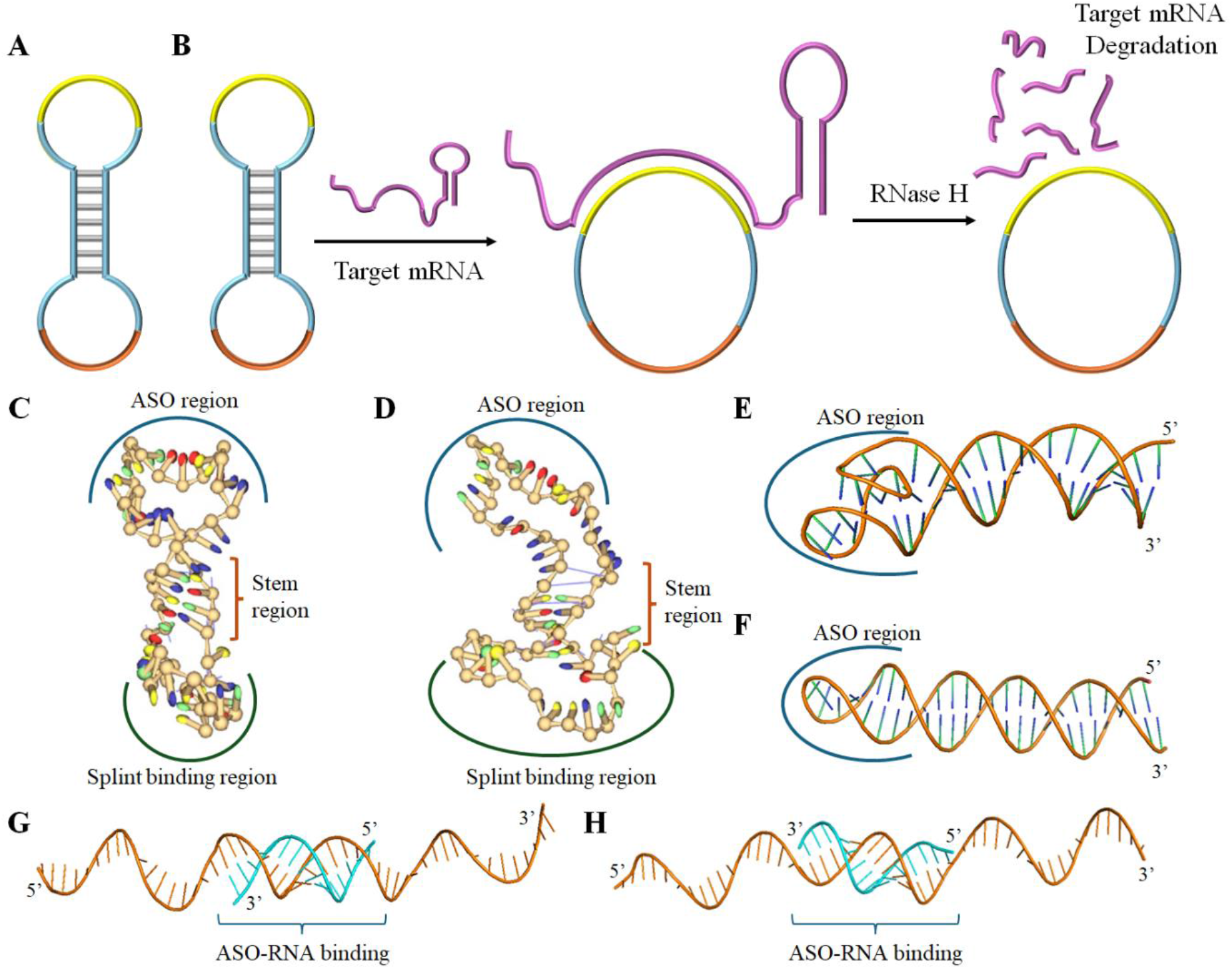
Structural design and proposed mechanism of action of NaFASO. A, Representative secondary structure of the NaFASO design, where the yellow, blue, and orange regions represent the ASO region, flexible stem, and the splint-padlock circularization domain. B, Mechanism of action of NaFASO—upon cellular entry, the ASO region (represented in yellow) hybridizes with the RNA target, destabilizing the stem region to support binding and subsequent RNase H-mediated target degradation, resulting in effective target knockdown. C and D, The oxView-predicted three-dimensional structures of NaFASO JINR1-1 and NaFASO JINR1-2, respectively. E and F, AlphaFold-predicted structures of precursors of NaFASO JINR1-1 (panel E) and NaFASO JINR1-2 with the ASO region highlighted. G and H, AlphaFold-based simulations illustrating RNA (blue)–NaFASO precursor interactions and stem structure-switching for NaFASO JINR1-1 and NaFASO JINR1-2, respectively.

We assigned the stem region a central role in promoting circularization. By juxtaposing the termini, the stem facilitated splint hybridization and ensured robust ligation. To prevent unintended interactions, we deliberately engineered the sequences outside the ASO region to minimize complementarity with mammalian nucleic acids, thereby reducing the risk of off-target binding. For optimal performance, we set the GC content of the stem at 50–60%. This balance provided sufficient thermodynamic stability to maintain stem integrity while allowing flexibility for transient opening (below). The design thereby enabled the ASO region to access its complementary target sequence without interference from other domains of the construct.

In addition, the NaFASO construct behaved analogously to a molecular beacon (Fig. 1B): the metastable 5 nt stem maintained a closed, stable conformation under physiological conditions, while target binding triggers structural opening to expose the ASO domain, thereby facilitating efficient hybridization. The structure-switching behavior was substantiated by AlphaFold-based simulations, which revealed structural change in NaFASO from the absence (Fig. 1E, F) to the presence (Fig. 1G, H) of the RNA target (blue). This architecture provided three key advantages. First, the ASO region remained fully accessible to its target with minimal structural hindrance from the rest of the construct or the stem itself. Second, non-specific interactions were suppressed enhancing both stability and knockdown specificity. Third, the two stem separated regions can be utilized to combinatorially incorporates independent functional domains (e.g., ASO, aptamers, DNAzymes etc) without affecting each other. Together, these design considerations established NaFASO as a rationally engineered, circular oligonucleotide nanostructure with tuneable secondary structure, optimized for nuclease resistance and precise gene silencing.

### Synthesis and characterization of NaFASO

To overcome the limitations of chemical synthesis, we designed the NaFASOs using splint padlock-based ligation where linear DNA precursors were subjected to 5′ phosphorylation, annealing with a splint, ligation, and subsequent exonuclease digestion (Fig. 2A). The efficiency of circularization was evaluated by loading the precursor, ligated, and exonuclease-digested samples onto a 12% denaturing urea PAGE. As shown in Fig. 2B and C, NaFASO JINR1-1 and JINR1-2 displayed a single band for their respective precursor (lane 2).

**Fig. 2.**
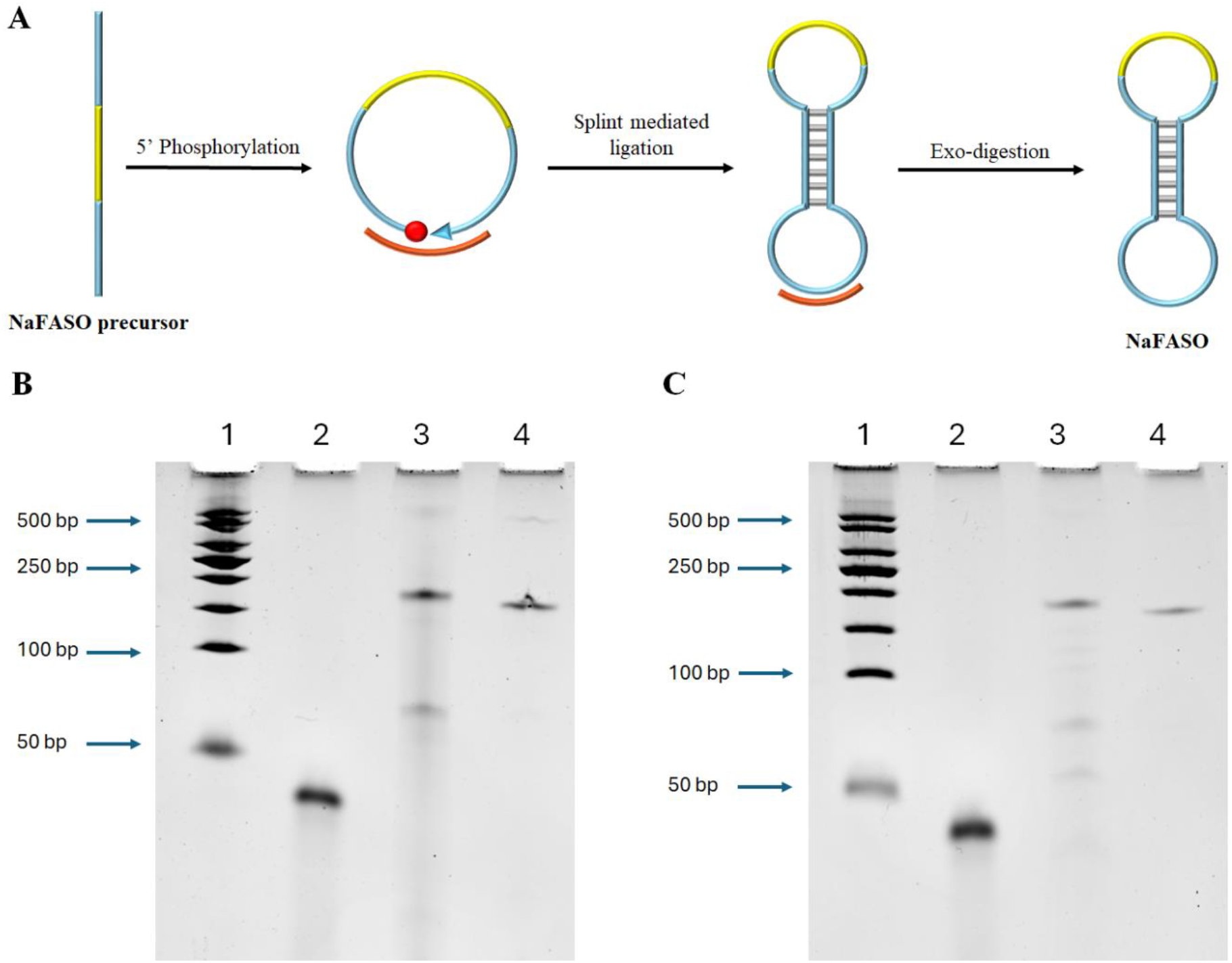
Synthesis and characterisation of NaFASOs. A, The schematic illustration of the synthesis of NaFASO. Linear DNA was phosphorylated, annealed with the splint, ligated, and exo-digested to remove linear DNA. B–C, Representative 12% denaturing urea PAGE showing NaFASO JINR1-1, and NaFASO JINR1-2 (respectively) characterisation after different synthesis steps. Lane 1, Ladder (50 bps), Lane 2, untreated control, Lane 3, Ligated sample, Lane 4, Exo digested sample (100 ng each).

In contrast, the ligated sample (lane 3) exhibited a slower-migrating band corresponding to circular DNA, along with additional faint bands and smearing indicative of incomplete ligation (Fig. 2B and C). The absence of the precursor band in the ligated sample confirmed successful end-to-end joining. Following exonuclease digestion (lane 4), only a single band corresponding to the circularized product was retained, verifying that all residual linear or unligated DNA was efficiently removed (Fig. 2B and C). In both cases, the presence of a distinct, slower-migrating band after ligation, along with its resistance to exonuclease digestion, validated the successful generation of nuclease- and serum-stable circular NaFASO constructs

### Stability of linear unmodified DNA v/s NaFASO

Therapeutic oligonucleotides typically contain modified sugar, phosphorothioated linkages, or both that confer stability against predominantly 3’-exonuclease class of endogenous nucleases^43^. To evaluate the nuclease resistance of NaFASO relative to unmodified linear DNA, we performed a comparative serum and nuclease stability assay using serum-spiked SVPD, one of the most potent existing exonucleases and a benchmark for testing oligonucleotide stability^44,45^ (Fig. 3A). As shown in Fig. 3B, unmodified linear DNA exhibited rapid degradation, with detectable band loss and smearing apparent within 5 min and more than 50% degradation by 30 min. In contrast, NaFASO (Fig. 3C) displayed remarkable stability, with distinct bands persisting without detectable smearing even after 90 min of incubation. Densitometric analysis (Fig. 3D) further confirmed that the band intensity of unmodified linear DNA (blue) decreased progressively over time, whereas NaFASO (orange) maintained a consistent signal throughout the entire incubation period. These findings establish that NaFASO circularization provided significant resistance against SVPD exonuclease-mediated degradation, underscoring its advantage over unmodified linear DNA constructs for applications requiring prolonged stability in biologically relevant environments. This resistance is significant in the context of antiviral therapy, as it ensures the sustained therapeutic effect over an extended period, thus enhancing the potential efficacy of NaFASO as a therapeutic agent.

**Fig. 3.**
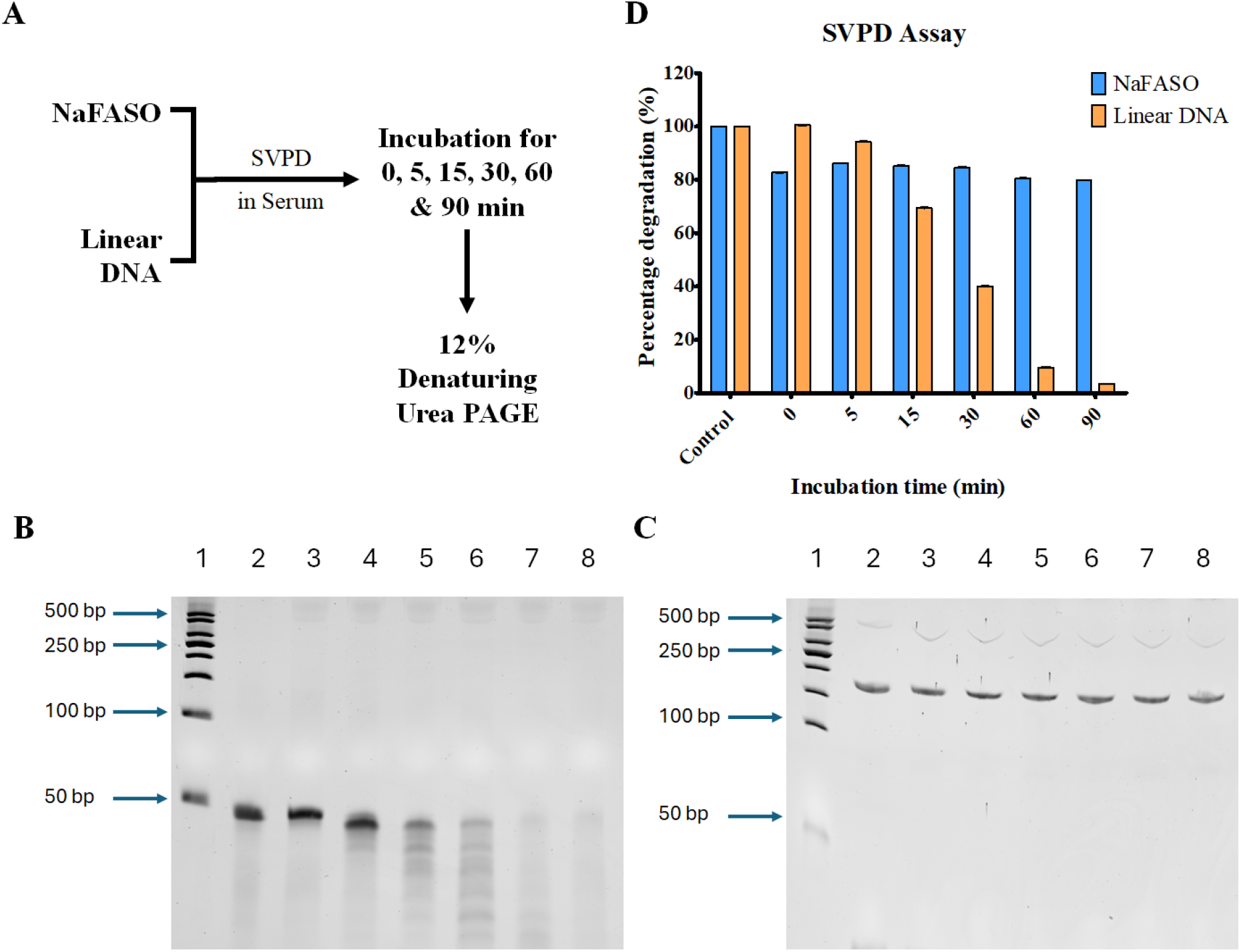
Serum stability assay of NaFASO versus linear precursors. A, Schematics of the SVPD assay, where samples were incubated with SVPD-spiked 10% FBS-containing serum for 0–90 min and analysed by 12% denaturing urea PAGE. B, Densitometric analysis showing degradation kinetics of linear DNA (orange) and NaFASO (blue). Error bars (n = 3) represent standard deviations). C–D, Representative 12% denaturing urea PAGE showing stability profiles of linear DNA (panel C), and NaFASO JINR1-1 (panel D) after SVPD treatment in serum. Lane 1, Ladder (50 bps), Lane 2, untreated control (250 ng), Lane 3-8, 0-, 5-, 15-, 30-, 60-, and 90-min samples respectively (250 ng each).

### NaFASO suppresses JINR1 expression and attenuates JEV-mediated cell death

First, we evaluated the comparative knockdown efficiency of NaFASO and LNA ASO against JINR1 in SHSY5Y cells. NaFASO displayed comparable knockdown efficiency to that of LNA ASO (Fig. S2), indicating that NaFASO achieves effective silencing without compromising performance. Next, we evaluated the impact of NaFASO on JINR1 expression during JEV infection and JEV-induced cell death. JEV infection significantly enhances JINR1 expression, resulting in a ∼5.14-fold increase in scrambled NaFASO–transfected cells compared to scrambled NaFASO–transfected mock-infected cells (Fig. 4B), consistent with previous reports by Tripathi et al^40^. Importantly, NaFASO-mediated JINR1 knockdown markedly attenuated this virus-induced JINR1 expression. During JEV infection, NaFASO JINR1-1 and NaFASO JINR1-2 reduce JINR1 expression to 2.2-fold and 2.70-fold, respectively, compared to scrambled NaFASO-transfected SH-SY5Y cells (Fig. 4B). These results demonstrate that NaFASO effectively reduces JEV-induced JINR1 expression. Notably, NaFASO JINR1-1 shows greater knockdown efficiency than NaFASO JINR1-2.

**Fig. 4.**
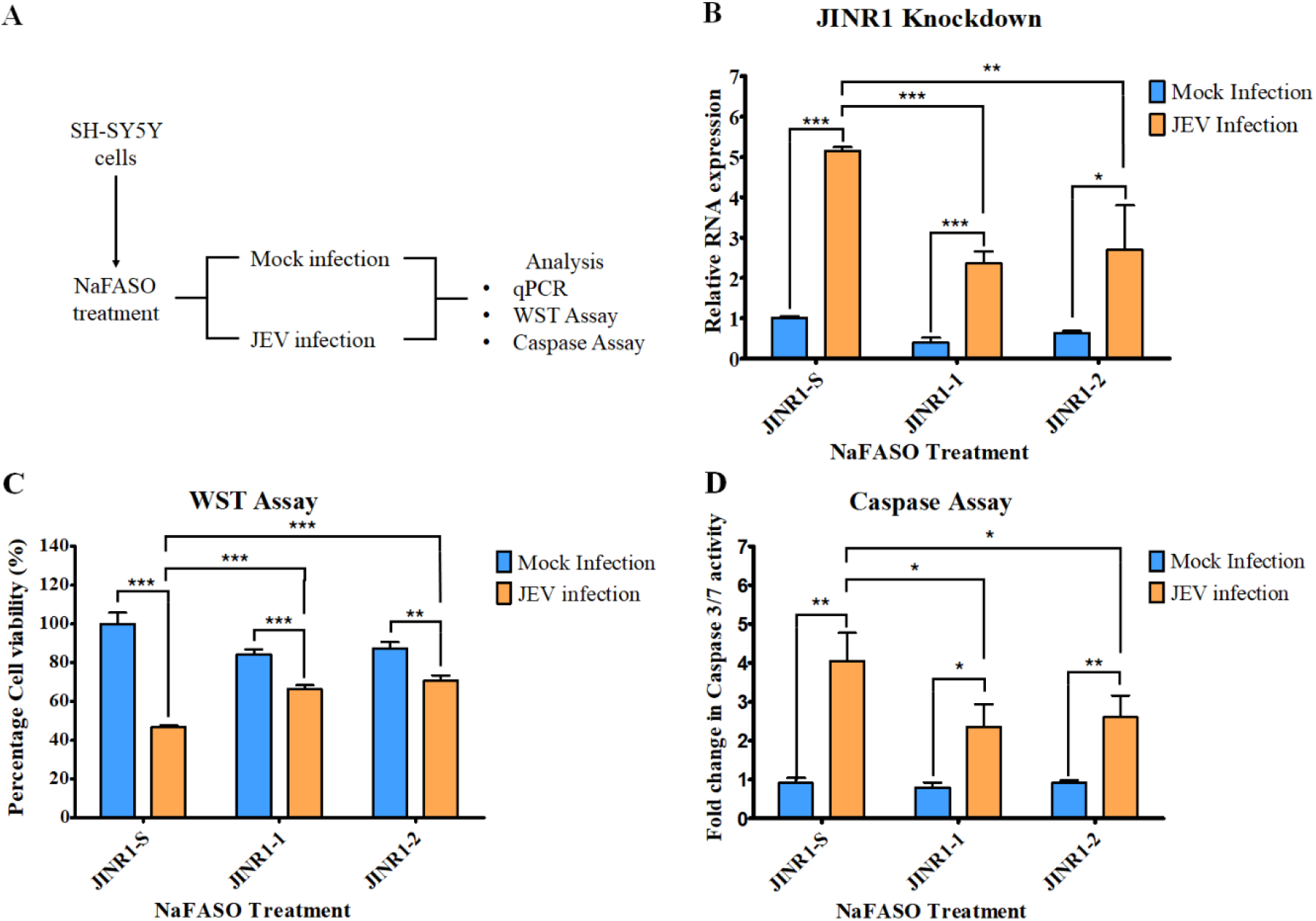
NaFASO-Mediated JINR1 knockdown modulates cell viability and apoptosis in JEV-Infected SH-SY5Y Cells. A, Schematic representation of the experimental workflow. SH-SY5Y cells reverse-transfected with scrambled NaFASO (control), NaFASO JINR1-1, or NaFASO JINR1-2, followed by JEV infection and subsequent analysis. B, qRT-PCR analysis showing relative JINR1 expression upon NaFASO treatment in Mock and JEV-infected cells. C, Cell proliferation assessed by WST-1 assay at 48 h post-infection (hpi) following JINR1 knockdown, presented as percentage of viable cells treated with scrambled NaFASO (control), NaFASO JINR1-1, or NaFASO JINR1-2. D, Apoptosis evaluated by measuring Caspase-3/7 activity at 48 hpi following JINR1 knockdown, represented as fold change in Caspase-3/7 activity in cells treated with scrambled NaFASO (control), NaFASO JINR1-1, or NaFASO JINR1-2. Error bars (n = 3) represents standard deviation. Statistical analysis performed using Student’s t-test. P < 0.05 (*), P < 0.01 (**), P < 0.001 (***)

To evaluate the impact of NaFASO-mediated JINR1 knockdown on JEV-induced cytotoxicity and apoptosis, cell proliferation and caspase-3/7 activity were measured using WST-1 reagent and caspase-3/7 assays, respectively. Cell proliferation analysis at 48 hpi revealed that, under mock-infected conditions, transfection with NaFASO JINR1-1 and NaFASO JINR1-2 resulted in only a modest reduction in viability (∼15% and ∼13%, respectively), indicating minimal intrinsic cytotoxicity associated with NaFASO treatment alone (Fig. 4C). In contrast, JEV infection markedly impaired cell viability, reducing proliferation to ∼47%, thereby confirming the pronounced cytopathic effect of viral infection. Notably, NaFASO-mediated JINR1 knockdown significantly attenuated this virus-induced loss of viability, with proliferation levels recovering to ∼66% and ∼71% in cells transfected with NaFASO JINR1-1 and NaFASO JINR1-2, respectively (Fig. 4C). These findings indicate that suppression of JINR1 partially restores cell viability during JEV infection. Consistent with the observed reduction in cell viability, JEV infection induced a pronounced apoptotic response, as evidenced by ∼5.14-fold increase in caspase-3/7 activity compared with mock-infected cells (Fig. 4D). Importantly, JINR1 knockdown markedly suppressed JEV-induced apoptosis. Cells transfected with NaFASO JINR1-1 exhibited an approximately 41.7% reduction in caspase 3/7 activity relative to JEV-infected cells transfected with scrambled NaFASO, while NaFASO JINR1-2 conferred a comparatively moderate reduction of ∼35.5% (Fig. 4D). Collectively, these results demonstrate that NaFASO-mediated suppression of *JINR1* alleviates JEV-induced cytotoxicity by both preserving cell viability and attenuating virus-driven apoptotic cell death, thereby underscoring *JINR1* as a potential host target for therapeutic intervention against JEV infection.

### Effect of NaFASO-mediated *JINR1* knockdown on JEV replication

Since NaFASO-mediated *JINR1* knockdown attenuated JEV-induced cell death, we evaluated the impact of NaFASO-mediated *JINR1* knockdown on JEV replication. We measured JEV RNA levels and the release of infectious virus particles upon NaFASO-mediated JINR1 depletion in SH-SY5Y cells. JEV RNA levels were measured by qRT-PCR, and a plaque assay was performed to determine viral titer, expressed as PFU/mL. Transfection with JINR1-targeting NaFASO construct significantly reduced JEV RNA levels compared to cells transfected with scrambled NaFASO (JINR1-S) (Fig. 5A). Specifically, NaFASO JINR1-1 led to a marked decrease in intracellular viral RNA abundance by 40% as compared to NaFASO JINR1-2 (27% reduction), indicating impaired viral replication. Consistent with these findings, JINR1-targeting NaFASO constructs displayed a significant decrease in viral titer corresponding to an approximate 51% reduction relative to JINR1-S–transfected cells (Fig. 5B). Similarly, NaFASO JINR1-2 significantly reduced JEV RNA levels and viral titers, representing a 35% reduction in titer (Fig. 5A and 5B). However, this effect was less pronounced than that observed with NaFASO JINR1-1, further supporting JINR1-1’s comparatively higher silencing efficiency. Collectively, these results demonstrate that NaFASO-mediated JINR1 knockdown restricts JEV replication in SH-SY5Y cells. The reduction in viral replication correlates with decreased caspase-3/7 activation and attenuated apoptotic cell death observed upon NaFASO treatment, further supporting a role for JINR1 in promoting efficient JEV replication and virus-induced cytopathology.

**Fig. 5.**
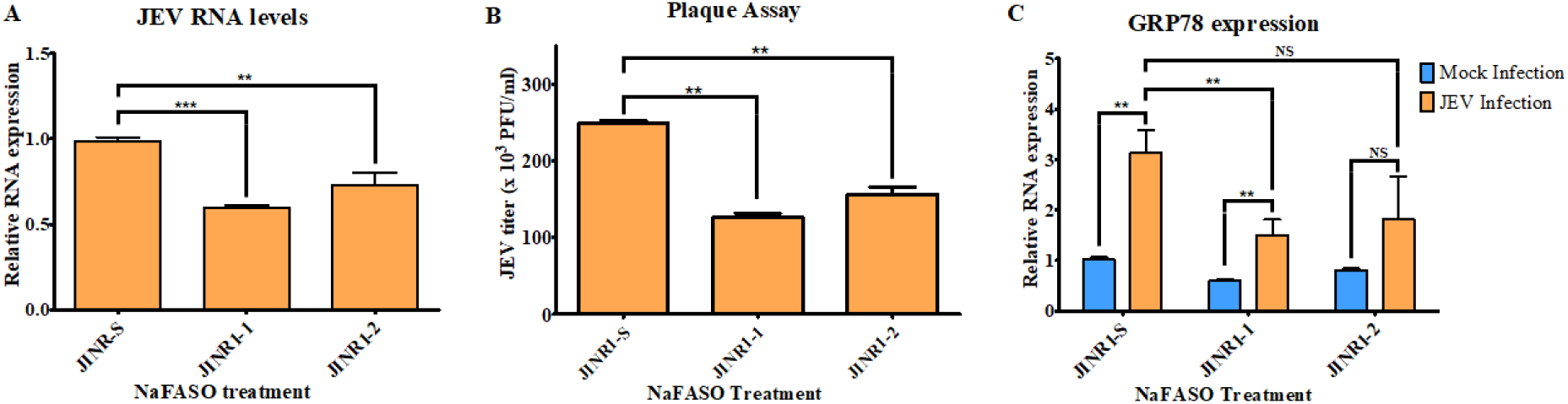
Effect of NaFASO-Mediated JINR1 knockdown on JEV replication and GRP78 expression. A, Quantitative real-time PCR (qRT-PCR) analysis of JEV RNA levels upon JINR1 silencing. B, Quantification of viral titers expressed as plaque-forming units per milliliter (PFU/mL), demonstrating the effect of JINR1 knockdown on JEV replication efficiency. C, Relative expression level of the downstream target gene GRP78 in cells treated with NaFASO constructs in the presence or absence of JEV infection. Error bars (n = 3) represents standard deviation. Statistical analysis performed using Student’s t-test. P < 0.05 (*), P < 0.01 (**), P < 0.001 (***)

GRP78 is known to promote JEV entry and replication in host cells and act as an important host factor for JEV pathogenesis ^40,41,46^. Moreover, the increase in GRP78 expression by JINR1 promotes virus replication^40^. Hence, we examined the effect of NaFASO-mediated JINR1 knockdown on GRP78 expression during JEV infection. SH-SY5Y cells were transfected with NaFASO JINR1-S, NaFASO JINR1-1, or NaFASO JINR1-2 and either mock-infected or infected with JEV, followed by qRT-PCR analysis of GRP78 expression. Under mock-infected conditions, NaFASO JINR1-1 and NaFASO JINR1-2 reduced basal GRP78 expression by approximately 40% and 20%, respectively, compared with scrambled NaFASO (JINR1-S) (Fig. 5C). NaFASO-mediated JINR1 knockdown significantly attenuated GRP78 expression to 1.49-fold (52% reduction relative to JINR1-S) and 1.81-fold (42% reduction) in NaFASO JINR1-1– and NaFASO JINR1-2–transfected cells, respectively (Fig. 5C). Consistent with previous observations, NaFASO JINR1-1 exhibited greater efficacy than NaFASO JINR1-2 in suppressing GRP78 expression (Fig. 5C). Collectively, these findings establish a functional link between NaFASO-mediated JINR1 knockdown, GRP78 suppression, and reduced JEV replication, highlighting NaFASO JINR1-1 as a consistently superior antiviral tool.

## Discussion

The alarming increase in JEV incidence across rural regions of South and South-east Asia represents a growing public health concern. As of today, there are no approved pathogen-directed antiviral therapies for JEV, aside from supportive management, which focuses on neurocritical care and seizure control ^44–46^. While multiple vaccine candidates, including inactivated and live-attenuated formulations, are approved, these approaches primarily serve as preventive measures rather than therapeutics, particularly for adult populations and individuals in remote settings^47,48^. While several small-molecule candidates show promising inhibition of JEV replication in cell culture and animal models, none have received regulatory approval ^49,50^.

We have shown that JINR1 is a critical regulator of orthoflavivirus replication and promotes neuronal cell death by directly interacting with RBM10 and NF-κB p65, thereby modulating the expression of GRP78 and other NF-κB target genes^40^. In addition, *JINR1* acts as a competing endogenous RNA (ceRNA) by sequestering *miR-216b-5p* and *miR-1-3p*, thereby increasing the expression of host factors GRP78 and DDX5, respectively, to facilitate JEV pathogenesis^41^. More recently, our findings in HNSCC cells revealed that JINR1 acts as a ceRNA for miR-1-3p and miR-216b-5p, resulting in the upregulation of Slug and GRP78, respectively^58^.

Beyond viral infection, JINR1 has been implicated in diverse pathological contexts. It is upregulated in OECC cells, where it promotes cellular proliferation and suppresses apoptosis by activating the PI3K/Akt signaling pathway via miR-1-3p^51^. JINR1 is also induced by TGF-β in Tenon’s fibroblasts and glaucoma tissues, promoting proliferation, migration, and autophagy via sequestration of miR-216b-5p^52^. In prostate cancer (PCa), elevated LINC01518 expression enhances CNKSR2 levels by sponging hsa-miR-320a, thereby contributing to tumor progression^53^. Collectively, these observations highlight the multifaceted regulatory functions of JINR1 across viral and non-viral disease settings, further supporting its potential as a therapeutic target.

ASOs, a powerful modality in modern medicine, offer a rational strategy for accessing otherwise “undruggable” targets such as lncRNAs with high precision compared to small molecules or biologics. The recent evolution of backbone chemistry has led to several ASO candidates being considered in clinical trials against lncRNAs implicated in both infectious and non-infectious diseases. Specifically, applications for silencing oncogenic lncRNAs, such as MALAT1 and HOTAIR, in solid tumors and hematologic malignancies have utilized gapmer ASOs ^47,54^. Similarly, targeting host immunomodulatory lncRNAs, such as lncRNA MIR99AHG, mitigated *Mycobacterium tuberculosis* by restoring antimicrobial effector functions ^55^.

Given the versatility of antisense platforms, antisense-mediated knockdown of lncRNA *JINR1* is thus anticipated to become an attractive option for attenuating JEV disease severity. Despite the transformative promise of ASOs, their widespread adoption has been restricted by challenges such as scalability issues, the limited viability of chemical synthesis beyond short (16-20 nt) oligonucleotides due to chemical modifications, and the generation of substantial amounts of hazardous waste. While alternative synthesis pathways involving biocatalytic or biosynthetic platforms circumvent these issues, they remain dependent on specialized enzymes and precursors, posing a bottleneck towards broadening the approaches for therapeutic ASO production. For instance, a gapmer ASO modality targeting *JINR1* in the JEV infection model demonstrated significant knockdown of *JINR1*, reduced downstream GRP78 levels, and decreased viral RNA levels^40^. However, this approach relied on LNA modality linear ASOs, which, although highly effective, are again limited by cost and scalability.

In the present study, we addressed these limitations by developing a biocatalytic synthesis method capable of generating a circular single-stranded DNA-based antisense construct, termed NaFASO, that integrates structural stability with functional specificity. The nanoconstruct was meticulously designed to comprise three distinct domains: a functional antisense region, a splint-binding region, and a stem region that spatially separates the two. To eliminate the requirement for backbone modification–mediated stabilization, a robust splint– padlock-based circularization strategy was optimized. Effective circularization effectively sealed the free 3′- and 5′-termini, thereby protecting NaFASO from exonuclease-mediated degradation and enhancing its stability under in vivo–relevant serum and SVPD nuclease conditions. As shown by oxView and AlphaFold simulation, the stem region plays a central role in regulating both the structural integrity and functional adaptability of the nanoconstruct. In addition to serving as a domain separator, the stem, in the absence of target RNA, adopts a stable secondary structure that facilitates efficient splint hybridization, enabling circularization. Upon target RNA binding, the metastable stem then undergoes controlled unzipping to enable effective ASO–target hybridization. This intrinsic structure-switching capability represents a defining feature of the NaFASO architecture and underlies its ability to balance stability with target accessibility.

Our experimental validation confirmed the feasibility and therapeutic promise of NaFASO. A comparative evaluation of LNA-modified ASOs (JINR1-1 and JINR1-2) and their corresponding NaFASO constructs demonstrated comparable knockdown efficiency under basal conditions (Fig. S2). This observation suggests that incorporation of the ASO sequence into the NaFASO nanoconstruct does not compromise silencing efficacy, supporting the potential of NaFASO as an effective alternative to conventional LNA-modified ASOs.In addition to withstanding the serum and nuclease degradation, functional assays involving NaFASO constructs further revealed that NaFASO exerted protective and anti-apoptotic effects. This was evidenced by improved cell viability and reduced caspase activation in both uninfected and JEV-infected cells. These findings prompted further investigation into the mechanistic consequences of NaFASO-mediated *JINR1* silencing during JEV infection.

Importantly, mechanistic analyses indicated that NaFASO effectively knocked down lncRNA *JINR1* expression in both contexts, resulting in a consequential decrease in GRP78 expression, a key host receptor that facilitates JEV entry. This downregulation was correlated with significant reductions in JEV RNA levels, thereby attenuating viral replication. The results were independently corroborated by plaque assays, which demonstrated a marked decrease in viral titre upon NaFASO treatment. Through the functional efficacy of NaFASO in JINR1 knockdown, our study validated the seamless integration of the antisense domain, the splint-padlock ligation domain, and the flexible structure-switching stem region. This demonstrated that the NaFASO constructs can overcome the length-related constraints of solid-phase synthesis. Furthermore, the synthesis was performed with “regular” enzymes (T4 PNK, T4 DNA ligase, exonucleases) routinely used in cloning and polymerase chain reaction clean-up assays^56^, thereby avoiding the need for specialized enzymes. The present work did not pursue the dosage determination of the NaFASO constructs and only validated the proof-of-concept therapeutic feasibility of enzyme-synthesized circular structure-switching ASOs.

Collectively, this work established NaFASOs as a novel, nuclease-resistant, serum-stable, therapeutically functional, yet scalable, cost-effective, and environmentally sustainable alternative to conventionally synthesized oligonucleotides. The NaFASO modality, for the first time (to the best of our knowledge), presents a circular antisense construct in which the gapmer antisense domain is effectively independent of the other functional regions within it. As a result, any domain of NaFASO that may attenuate the antisense region’s efficacy was thus separated by the metastable stem analogous to that in a molecular beacon. This work would also be the first (to the best of our knowledge) demonstration of a biocatalytically synthesized ASO that successfully targeted a host lncRNA in the JEV model. The broader implication of this work is that biocatalytic strategies for ASO synthesis using regular enzymes can achieve meaningful biological outcomes, thereby setting the stage for next-generation ASO therapeutics. Accordingly, future studies will assess the dose-response of the NaFASOs and identify optimal conditions to achieve maximal therapeutic efficacy without introducing toxicity. Furthermore, we will study the placement and subsequent effects of additional functional nucleic acid entities such as aptamers in stem-separated regions to further explore cell/tissue selectivity and specificity in various disease models.

## Conclusion

In summary, this proof-of-concept study demonstrates the biocatalytic synthesis of NaFASO, a novel circular ASO construct (NaFASO) that exhibits stability, specificity, and therapeutic efficacy against a host lncRNA as a therapeutic target during JEV infection. By targeting lncRNA *JINR1*, NaFASO not only disrupted viral replication but also protected the host cell against virus-induced apoptosis, underscoring the dual functional potential of this design. Compared to conventionally synthesized ASOs, NaFASO represents a sustainable, cost-effective alternative that avoids many of the drawbacks associated with backbone modifications and environmentally hazardous chemical processes while offering structural and design flexibility. The success of this strategy illustrates that enzyme-mediated approaches can yield oligonucleotide therapeutics of high fidelity and biological relevance. Looking forward, expanding the substrate tolerance of enzymatic platforms will be essential for broader clinical translation. Future efforts would thus focus on dosage optimization, preclinical studies, adapting this methodology to target a broader range of RNA sequences, expanding its applications to oncology and genetic disorders, and developing scalable production pipelines. Together, these advances have the potential to reshape the trajectory of ASO therapeutics toward more accessible and environmentally responsible medicine.

## Author contributions

CS, SS, VS, and SG designed the study. CS, SS, and DS performed the experiments. CS, SS, and DS performed the data analysis. CS, SS, VS, and SG wrote the manuscript. All authors edited and approved the final manuscript.

## Data Availability

The data for this manuscript is available upon reasonable request.

## Acknowledgement

This work was supported by the DST-Anusandhan National Research Foundation (ANRF)-CRG/2023/000822 to VS, and Mahindra University internal funding. The authors are grateful to Prof. Rajinder S. Chauhan (Dean, Centre for Life Sciences) and Prof Yajulu Medury (Vice Chancellor, Mahindra University) for their constant scholarly encouragement. We acknowledge the use of the grammar correction tool “Grammarly” in editing the manuscript. SS is supported by CSIR SRF fellowship (File no. 09/1026(23714/2025-EMR-I)

## Electronic Supplementary Information (ESI)

Oligonucleotide sequences, NaFASO secondary structures, and plaque assay representative images.

## References

(1) Roberts, T. C.; Langer, R.; Wood, M. J. A. Advances in Oligonucleotide Drug Delivery. Nat. Rev. Drug Discov. 2020, 19 (10), 673–694. 10.1038/s41573-020-0075-7.

(2) Collotta, D.; Bertocchi, I.; Chiapello, E.; Collino, M. Antisense Oligonucleotides: A Novel Frontier in Pharmacological Strategy. Front. Pharmacol. 2023, 14. 10.3389/fphar.2023.1304342.

(3) Shevelev, A.; Pozdniakova, N.; Generalov, E.; Tarasova, O. SiRNA Therapeutics for the Treatment of Hereditary Diseases and Other Conditions: A Review. Int. J. Mol. Sci. 2025, 26 (17), 8651. 10.3390/ijms26178651.

(4) Trivedi, J.; Yasir, M.; Maurya, R. K.; Tripathi, A. S. Aptamer-Based Theranostics in Oncology: Design Strategies and Limitations. BIO Integr. 2024, 5 (1). 10.15212/bioi-2024-0002.

(5) Xie, X.; Yu, T.; Li, X.; Zhang, N.; Foster, L. J.; Peng, C.; Huang, W.; He, G. Recent Advances in Targeting the “Undruggable” Proteins: From Drug Discovery to Clinical Trials. Signal Transduct. Target. Ther. 2023, 8 (1), 335. 10.1038/s41392-023-01589-z.

(6) Finkel, R. S.; Mercuri, E.; Darras, B. T.; Connolly, A. M.; Kuntz, N. L.; Kirschner, J.; Chiriboga, C. A.; Saito, K.; Servais, L.; Tizzano, E.; Topaloglu, H.; Tulinius, M.; Montes, J.; Glanzman, A. M.; Bishop, K.; Zhong, Z. J.; Gheuens, S.; Bennett, C. F.; Schneider, E.; Farwell, W.; Vivo, D. C. D. Nusinersen versus Sham Control in Infantile-Onset Spinal Muscular Atrophy. N. Engl. J. Med. 2017, 377 (18), 1723–1732. 10.1056/NEJMoa1702752.

(7) Mitelman, O.; Abdel-Hamid, H. Z.; Byrne, B. J.; Connolly, A. M.; Heydemann, P.; Proud, C.; Shieh, P. B.; Wagner, K. R.; Dugar, A.; Santra, S.; Signorovitch, J.; Goemans, N.; McDonald, C. M.; Mercuri, E.; Mendell, J. R. A Combined Prospective and Retrospective Comparison of Long-Term Functional Outcomes Suggests Delayed Loss of Ambulation and Pulmonary Decline with Long-Term Eteplirsen Treatment. J. Neuromuscul. Dis. 9 (1), 39–52. 10.3233/JND-210665.

(8) Epple, S.; El-Sagheer, A. H.; Brown, T. Artificial Nucleic Acid Backbones and Their Applications in Therapeutics, Synthetic Biology and Biotechnology. Emerg. Top. Life Sci. 2021, 5 (5), 691–697. 10.1042/ETLS20210169.

(9) Liu, C.; Cozens, C.; Jaziri, F.; Rozenski, J.; Maréchal, A.; Dumbre, S.; Pezo, V.; Marlière, P.; Pinheiro, V. B.; Groaz, E.; Herdewijn, P. Phosphonomethyl Oligonucleotides as Backbone-Modified Artificial Genetic Polymers. J. Am. Chem. Soc. 2018, 140 (21), 6690–6699. 10.1021/jacs.8b03447.

(10) Full article: Antisense oligonucleotides: absorption, distribution, metabolism, and excretion. https://www.tandfonline.com/doi/10.1080/17425255.2021.1992382?url_ver=Z39.88-2003&rfr_id=ori:rid:crossref.org&rfr_dat=cr_pub%20%200pubmed (accessed 2025-11-06).

(11) Hammond, S. M.; Aartsma-Rus, A.; Alves, S.; Borgos, S. E.; Buijsen, R. A. M.; Collin, R. W. J.; Covello, G.; Denti, M. A.; Desviat, L. R.; Echevarría, L.; Foged, C.; Gaina, G.; Garanto, A.; Goyenvalle, A. T.; Guzowska, M.; Holodnuka, I.; Jones, D. R.; Krause, S.; Lehto, T.; Montolio, M.; Van Roon-Mom, W.; Arechavala-Gomeza, V. Delivery of Oligonucleotide-based Therapeutics: Challenges and Opportunities. EMBO Mol. Med. 2021, 13 (4), e13243. 10.15252/emmm.202013243.

(12) Andrews, B. I.; Antia, F. D.; Brueggemeier, S. B.; Diorazio, L. J.; Koenig, S. G.; Kopach, M. E.; Lee, H.; Olbrich, M.; Watson, A. L. Sustainability Challenges and Opportunities in Oligonucleotide Manufacturing. J. Org. Chem. 2021, 86 (1), 49–61. 10.1021/acs.joc.0c02291.

(13) Herkt, M.; Thum, T. Pharmacokinetics and Proceedings in Clinical Application of Nucleic Acid Therapeutics. Mol. Ther. 2021, 29 (2), 521–539. 10.1016/j.ymthe.2020.11.008.

(14) Takakusa, H.; Iwazaki, N.; Nishikawa, M.; Yoshida, T.; Obika, S.; Inoue, T. Drug Metabolism and Pharmacokinetics of Antisense Oligonucleotide Therapeutics: Typical Profiles, Evaluation Approaches, and Points to Consider Compared with Small Molecule Drugs. Nucleic Acid Ther. 2023, 33 (2), 83–94. 10.1089/nat.2022.0054.

(15) Kumari, A.; Kaur, A.; Aggarwal, G. The Emerging Potential of SiRNA Nanotherapeutics in Treatment of Arthritis. Asian J. Pharm. Sci. 2023, 18 (5), 100845. 10.1016/j.ajps.2023.100845.

(16) Wang, Y.; Wu, L.; Wang, P.; Lv, C.; Yang, Z.; Tang, X. Manipulation of Gene Expression in Zebrafish Using Caged Circular Morpholino Oligomers. Nucleic Acids Res. 2012, 40 (21), 11155–11162. 10.1093/nar/gks840.

(17) Pandey, E.; Harris, E. N. Chloroquine and Cytosolic Galectins Affect Endosomal Escape of Antisense Oligonucleotides after Stabilin-Mediated Endocytosis. Mol. Ther. Nucleic Acids 2023, 33, 430–443. 10.1016/j.omtn.2023.07.019.

(18) Griepenburg, J. C.; Rapp, T. L.; Carroll, P. J.; Eberwine, J.; Dmochowski, I. J. Ruthenium-Caged Antisense Morpholinos for Regulating Gene Expression in Zebrafish Embryos. Chem. Sci. 2015, 6 (4), 2342–2346. 10.1039/C4SC03990D.

(19) Yang, L.; von Trentini, D.; Kim, H.; Sul, J.-Y.; Eberwine, J. H.; Dmochowski, I. J. Photoactivatable Circular Caged Oligonucleotides for Transcriptome In Vivo Analysis (TIVA). ChemPhotoChem 2021, 5 (10), 940–946. 10.1002/cptc.202100098.

(20) Jahns, H.; Degaonkar, R.; Podbevsek, P.; Gupta, S.; Bisbe, A.; Aluri, K.; Szeto, J.; Kumar, P.; LeBlanc, S.; Racie, T.; Brown, C. R.; Castoreno, A.; Guenther, D. C.; Jadhav, V.; Maier, M. A.; Plavec, J.; Egli, M.; Manoharan, M.; Zlatev, I. Small Circular Interfering RNAs (SciRNAs) as a Potent Therapeutic Platform for Gene-Silencing. Nucleic Acids Res. 2021, 49 (18), 10250–10264. 10.1093/nar/gkab724.

(21) Blumenfeld, M.; Brandys, P.; D’auriol, L.; Vasseur, M. Closed Sense and Antisense Oligonucleotides and Uses Thereof. WO1992019732A1, November 12, 1992. https://patents.google.com/patent/WO1992019732A1/en (accessed 2025-11-11).

(22) He, W.; Zhang, X.; Zou, Y.; Li, J.; Chang, L.; He, Y.-C.; Jin, Q.; Ye, J. Effective Synthesis of CircRNA via a Thermostable T7 RNA Polymerase Variant as the Catalyst. Front. Bioeng. Biotechnol. 2024, 12, 1356354. 10.3389/fbioe.2024.1356354.

(23) He, A. T.; Liu, J.; Li, F.; Yang, B. B. Targeting Circular RNAs as a Therapeutic Approach: Current Strategies and Challenges. Signal Transduct. Target. Ther. 2021, 6 (1), 185. 10.1038/s41392-021-00569-5.

(24) Nielsen, A. F.; Bindereif, A.; Bozzoni, I.; Hanan, M.; Hansen, T. B.; Irimia, M.; Kadener, S.; Kristensen, L. S.; Legnini, I.; Morlando, M.; Jarlstad Olesen, M. T.; Pasterkamp, R. J.; Preibisch, S.; Rajewsky, N.; Suenkel, C.; Kjems, J. Best Practice Standards for Circular RNA Research. Nat. Methods 2022, 19 (10), 1208–1220. 10.1038/s41592-022-01487-2.

(25) Kristensen, L. S.; Andersen, M. S.; Stagsted, L. V. W.; Ebbesen, K. K.; Hansen, T. B.; Kjems, J. The Biogenesis, Biology and Characterization of Circular RNAs. Nat. Rev. Genet. 2019, 20 (11), 675–691. 10.1038/s41576-019-0158-7.

(26) Tandale, B. V.; Deshmukh, P. S.; Tomar, S. J.; Narang, R.; Qazi, M. S.; Goteti Venkata, P.; Jain, M.; Jain, D.; Guduru, V. K.; Jain, J.; Gosavi, R. V.; Valupadas, C. S.; Deshmukh, P. R.; Raut, A. V.; Narlawar, U. W.; Jha, P. K.; Bondre, V. P.; Sapkal, G. N.; Damle, R. G.; Khude, P. M.; Niswade, A. K.; Talapalliwar, M.; Rathod, P.; Balla, P. S.; Muttineni, P. K.; Kalepally Janakiram, K. K.; Rajderkar, S. S. Incidence of Japanese Encephalitis and Acute Encephalitis Syndrome Hospitalizations in the Medium-Endemic Region in Central India. J. Epidemiol. Glob. Health 2023, 13 (2), 173–179. 10.1007/s44197-023-00110-7.

(27) Ashraf, U.; Ding, Z.; Deng, S.; Ye, J.; Cao, S.; Chen, Z. Pathogenicity and Virulence of Japanese Encephalitis Virus: Neuroinflammation and Neuronal Cell Damage. Virulence 2021, 12 (1), 968–980. 10.1080/21505594.2021.1899674.

(28) Cheng, Y.; Minh, N. T.; Minh, Q. T.; Khandelwal, S.; Clapham, H. E. Estimates of Japanese Encephalitis Mortality and Morbidity: A Systematic Review and Modeling Analysis. PLoS Negl. Trop. Dis. 2022, 16 (5), e0010361. 10.1371/journal.pntd.0010361.

(29) Mulvey, P.; Duong, V.; Boyer, S.; Burgess, G.; Williams, D. T.; Dussart, P.; Horwood, P. F. The Ecology and Evolution of Japanese Encephalitis Virus. Pathogens 2021, 10 (12), 1534. 10.3390/pathogens10121534.

(30) Chen, H.-Y.; Yang, C.-Y.; Hsieh, C.-Y.; Yeh, C.-Y.; Chen, C.-C.; Chen, Y.-C.; Lai, C.-C.; Harris, R. C.; Ou, H.-T.; Ko, N.-Y.; Ko, W.-C. Long-Term Neurological and Healthcare Burden of Adults with Japanese Encephalitis: A Nationwide Study 2000-2015. PLoS Negl. Trop. Dis. 2021, 15 (9), e0009703. 10.1371/journal.pntd.0009703.

(31) Duerlund, L. S.; Nielsen, H.; Bodilsen, J. Current Epidemiology of Infectious Encephalitis: A Narrative Review. Clin. Microbiol. Infect. 2025, 31 (4), 515–521. 10.1016/j.cmi.2024.12.025.

(32) Epidemiological Profile of Acute Viral Encephalitis | Indian Journal of Pediatrics. https://link.springer.com/article/10.1007/s12098-017-2481-3 (accessed 2025-11-12).

(33) Tandale, B. V.; Deshmukh, P. S.; Narang, R.; Qazi, M. S.; Padmaja, G. V.; Deshmukh, P. R.; Raut, A. V.; Narlawar, U. W.; Jha, P. K.; Rajderkar, S. S.; Group, J. E. E. S. Coverage of Japanese Encephalitis Routine Vaccination among Children in Central India. J. Med. Virol. 2023, 95 (1), e28155. 10.1002/jmv.28155.

(34) Rustagi, R.; Basu, S.; Garg, S. Japanese Encephalitis: Strategies for Prevention and Control in India. Indian J. Med. Spec. 2019, 10 (1), 12. 10.4103/INJMS.INJMS_22_18.

(35) Joe, S.; Salam, A. A. A.; Neogi, U. N N. B.; Mudgal, P. P. Antiviral Drug Research for Japanese Encephalitis: An Updated Review. Pharmacol. Rep. 2022, 74 (2), 273–296. 10.1007/s43440-022-00355-2.

(36) Letson, G. W.; Marfin, A. A.; Mooney, J.; Minh, H. V.; Hills, S. L.; Team, the J. V. G. I. A. Impact of Vaccination against Japanese Encephalitis in Endemic Countries. PLoS Negl. Trop. Dis. 2024, 18 (9), e0012390. 10.1371/journal.pntd.0012390.

(37) Vannice, K. S.; Hills, S. L.; Schwartz, L. M.; Barrett, A. D.; Heffelfinger, J.; Hombach, J.; Letson, G. W.; Solomon, T.; Marfin, A. A. The Future of Japanese Encephalitis Vaccination: Expert Recommendations for Achieving and Maintaining Optimal JE Control. Npj Vaccines 2021, 6 (1), 82. 10.1038/s41541-021-00338-z.

(38) Li, Y.; Zhang, H.; Zhu, B.; Ashraf, U.; Chen, Z.; Xu, Q.; Zhou, D.; Zheng, B.; Song, Y.; Chen, H.; Ye, J.; Cao, S. Microarray Analysis Identifies the Potential Role of Long Non-Coding RNA in Regulating Neuroinflammation during Japanese Encephalitis Virus Infection. Front. Immunol. 2017, 8. 10.3389/fimmu.2017.01237.

(39) Zhou, X.; Yuan, Q.; Zhang, C.; Dai, Z.; Du, C.; Wang, H.; Li, X.; Yang, S.; Zhao, A. Inhibition of Japanese Encephalitis Virus Proliferation by Long Non-Coding RNA SUSAJ1 in PK-15 Cells. Virol. J. 2021, 18 (1), 29. 10.1186/s12985-021-01492-5.

(40) Tripathi, S.; Sengar, S.; Shree, B.; Mohapatra, S.; Basu, A.; Sharma, V. An RBM10 and NF-KB Interacting Host LncRNA Promotes JEV Replication and Neuronal Cell Death. J. Virol. 2023, 97 (12), e0118323. 10.1128/jvi.01183-23.

(41) Tripathi, S.; Sengar, S.; Basu, A.; Sharma, V. LncRNA JINR1 Regulates MiR-216b-5p/GRP78 and MiR-1-3p/DDX5 Axis to Promote JEV Infection and Cell Death. J. Virol. 2025, 99 (5), e00066–25. 10.1128/jvi.00066-25.

(42) Bohlin, J.; Matthies, M.; Poppleton, E.; Procyk, J.; Mallya, A.; Yan, H.; Šulc, P. Design and Simulation of DNA, RNA and Hybrid Protein–Nucleic Acid Nanostructures with OxView. Nat. Protoc. 2022, 17 (8), 1762–1788. 10.1038/s41596-022-00688-5.

(43) Christmann, M.; Tomicic, M. T.; Aasland, D.; Berdelle, N.; Kaina, B. Three Prime Exonuclease I (TREX1) Is Fos/AP-1 Regulated by Genotoxic Stress and Protects against Ultraviolet Light and Benzo(a)Pyrene-Induced DNA Damage. Nucleic Acids Res. 2010, 38 (19), 6418–6432. 10.1093/nar/gkq455.

(44) Lovrić, J.; Yan, J.; Li, X.; Karlsborn, T.; Bood, M.; Dahlén, A.; Hilgendorf, C.; Bhatt, D. K. In Vitro Structure–Activity Relationship Stability Study of Antisense Oligonucleotide Therapeutics Using Biological Matrices and Nucleases. Pharmacol. Res. Perspect. 2025, 13 (3), e70096. 10.1002/prp2.70096.

(45) Monia, B. P.; Johnston, J. F.; Sasmor, H.; Cummins, L. L. Nuclease Resistance and Antisense Activity of Modified Oligonucleotides Targeted to Ha-Ras*. J. Biol. Chem. 1996, 271 (24), 14533–14540. 10.1074/jbc.271.24.14533.

(46) Nain, M.; Mukherjee, S.; Karmakar, S. P.; Paton, A. W.; Paton, J. C.; Abdin, M. Z.; Basu, A.; Kalia, M.; Vrati, S. GRP78 Is an Important Host Factor for Japanese Encephalitis Virus Entry and Replication in Mammalian Cells. J. Virol. 2017, 91 (6), 10.1128/jvi.02274-16. http://doi.org//10.1128/jvi.02274-16.

(47) Lu, J.; Guo, J.; Liu, J.; Mao, X.; Xu, K. Long Non-Coding RNA MALAT1: A Key Player in Liver Diseases. Front. Med. 2022, 8. 10.3389/fmed.2021.734643.

(48) Yun, S.-I.; Lee, Y.-M. Japanese Encephalitis: The Virus and Vaccines. Hum. Vaccines Immunother. 2014, 10 (2), 263–279. 10.4161/hv.26902.

(49) Yin, C.; Yang, P.; Xiao, Q.; Sun, P.; Zhang, X.; Zhao, J.; Hu, X.; Shan, C. Novel Antiviral Discoveries for Japanese Encephalitis Virus Infections through Reporter Virus-Based High-Throughput Screening. J. Med. Virol. 2024, 96 (1), e29382. 10.1002/jmv.29382.

(50) Guo, J.; Jia, X.; Liu, Y.; Wang, S.; Cao, J.; Zhang, B.; Xiao, G.; Wang, W. Screening of Natural Extracts for Inhibitors against Japanese Encephalitis Virus Infection. Antimicrob. Agents Chemother. 2020, 64 (3), 10.1128/aac.02373-19. http://doi.org/10.1128/aac.02373-19.

(51) Zhang, D.; Zhang, H.; Wang, X.; Hu, B.; Zhang, F.; Wei, H.; Li, L. LINC01518 Knockdown Inhibits Tumorigenicity by Suppression of PIK3CA/Akt Pathway in Oesophageal Squamous Cell Carcinoma. Artif. Cells Nanomedicine Biotechnol. 2019, 47 (1), 4284–4292. 10.1080/21691401.2019.1699815.

(52) Kong, N.; Bao, Y.; Zhao, H.; Kang, X.; Tai, X.; Chen, X.; Guo, W.; Shen, Y. Long Noncoding RNA LINC01518 Modulates Proliferation and Migration in TGF-B1-Treated Human Tenon Capsule Fibroblast Cells Through the Regulation of Hsa-MiR-216b-5p. Neuromolecular Med. 2022, 24 (2), 88–96. 10.1007/s12017-021-08662-2.

(53) Chen, G.; Chen, Z. LINC01518 Predicts Poor Prognosis of Prostate Cancer and Promotes Its Progression by Regulating Hsa-MiR-320a/CNKSR2 Axis. Discov. Oncol. 2024, 15 (1), 576. 10.1007/s12672-024-01458-3.

(54) Aiello, A.; Bacci, L.; Re, A.; Ripoli, C.; Pierconti, F.; Pinto, F.; Masetti, R.; Grassi, C.; Gaetano, C.; Bassi, P. F.; Pontecorvi, A.; Nanni, S.; Farsetti, A. MALAT1 and HOTAIR Long Non-Coding RNAs Play Opposite Role in Estrogen-Mediated Transcriptional Regulation in Prostate Cancer Cells. Sci. Rep. 2016, 6 (1), 38414. 10.1038/srep38414.

(55) Gcanga, L.; Tamgue, O.; Ozturk, M.; Pillay, S.; Jacobs, R.; Chia, J. E.; Mbandi, S. K.; Davids, M.; Dheda, K.; Schmeier, S.; Alam, T.; Roy, S.; Suzuki, H.; Brombacher, F.; Guler, R. Host-Directed Targeting of LincRNA-MIR99AHG Suppresses Intracellular Growth of Mycobacterium Tuberculosis. Nucleic Acid Ther. 2022, 32 (5), 421–437. 10.1089/nat.2022.0009.

(56) Watanabe, K.; Emoto, N.; Sunohara, M.; Kawakami, M.; Kage, H.; Nagase, T.; Ohishi, N.; Takai, D. Treatment of PCR Products with Exonuclease I and Heat-Labile Alkaline Phosphatase Improves the Visibility of Combined Bisulfite Restriction Analysis. Biochem. Biophys. Res. Commun. 2010, 399 (3), 422–424. 10.1016/j.bbrc.2010.07.093.

