## Supplemental Data for "Elucidating the effect of a rationally designed nanostructured form-switching ASO (NaFASO) for targeting long non-coding RNA to alleviate Japanese encephalitis virus infection"

#Equal contribution as co-first authors

**Supplementary Table S1:** List of sequences of various oligonucleotides used in different experiments

| **Name** | **Sequence (5'-3') (italics: splint binding region; underlined: antisense region)** |
|---|---|
| NaFASO JINR1-1 * | *GACCTGCTA*GACTGAAAATACAGCGCCTTTGGTAAAAACAGTC*CTGGACGAG* |
| NaFASO JINR1-2 ** | *GACCTGCTA*GACTGAAAAGAGGGAGAACAACGTTAAAACAGTC*CTGGACGAG* |
| NaFASO scrambled JINR1-S | *GACCTGCTA*GACTGAAAACGGAAAGAGTGCGATAAAACAGTC*CTGGACGAG* |
| LNA JINR1-1 | TACAGCGCCTTTGGTA |
| LNA JINR1-2 | GAGGGAGAACAACGTT |
| Splint | TAGCAGGTCCTCGTCCAG |
| JINR1 fwd primer | CAGTGACGGAACAGTACCAG |
| JINR1 rev primer | TCACAAACATCCCGCTCT |
| GRP78 fwd primer | CTGTCCAGGCTGGTGTGCTCT |
| GRP78 rev primer | CTTGGTAGGCACCACTGTGTTC |
| JEV fwd primer | GAGCTTGTTGGACGGCAGAG |
| JEV rev primer | CACGGCGTCGATGAGTGTTC |

*, ** Antisense region sequences obtained from Tripathi, S.; Sengar, S.; Shree, B.; Mohapatra, S.; Basu, A.; Sharma, V. An RBM10 and NF-KB Interacting Host LncRNA Promotes JEV Replication and Neuronal Cell Death. J. Virol. 2023, 97 (12), e0118323.

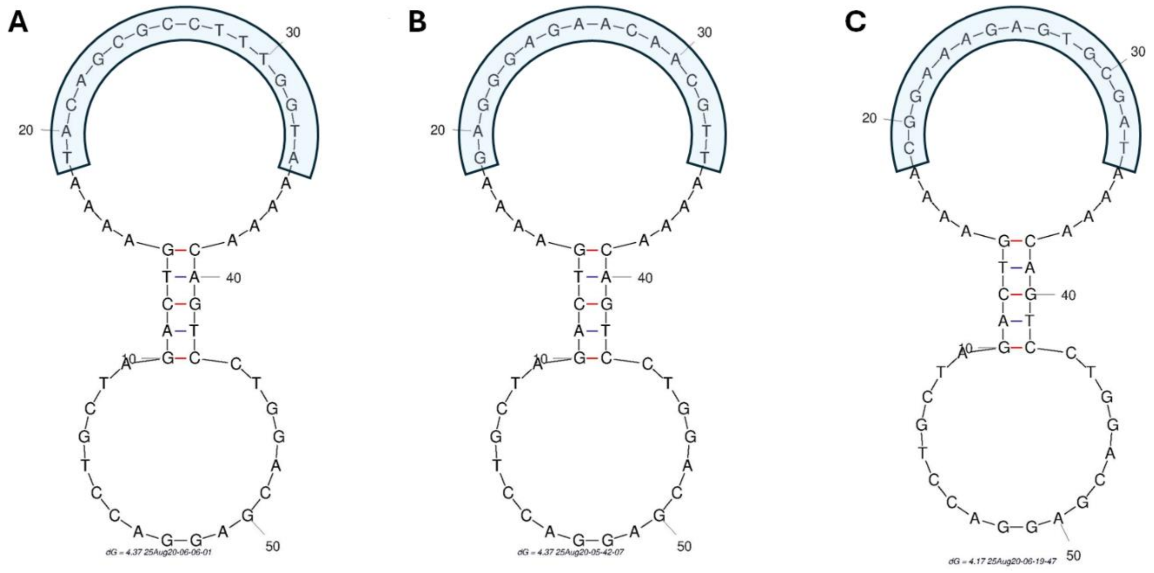


**Fig. S1** Secondary structure of NaFASO. A, mFold image of NaFASO JINR1-1 sequence with antisense oligonucleotide (ASO) sequence highlighted in blue. B, mFold image of NaFASO JINR1-2 sequence with ASO sequence highlighted in blue. C, mFold image of NaFASO scrambled JINR1-S with the scrambled ASO sequence highlighted in blue.

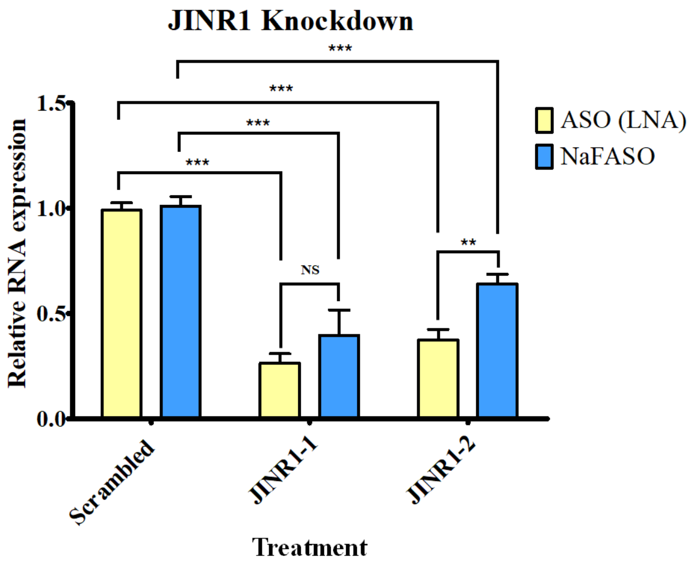


**Fig. S2** Comparative Efficiency of NaFASO and LNA-ASO–Mediated JINR1 Knockdown in SH-SY5Y Cells. Bar graph showing comparative analysis of NaFASO- and LNA-ASO–mediated JINR1 knockdown under basal (uninfected) conditions in SH-SY5Y cells, assessing the relative silencing efficiency of the two antisense strategies. Error bars (n = 3) represent standard deviation (SD). Statistical significance determined using Student's t-test. $P < 0.05$ (*), $P < 0.01$ (**), $P < 0.001$ (***); NS, not significant.

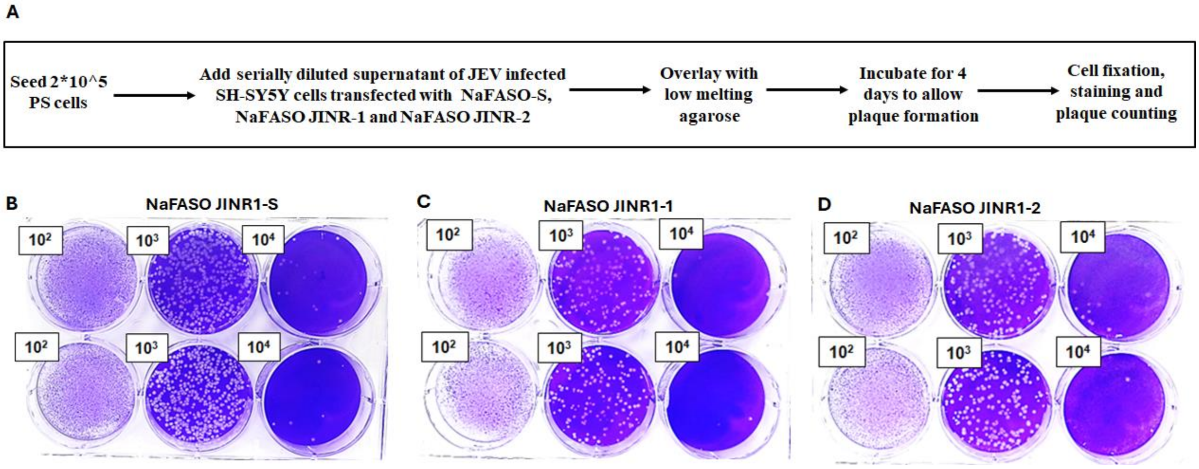


Fig. S3: Plaque assay workflow and representative results from different samples. A. Schematic representation of the experimental workflow used for the JEV plaque assay. B-D, Representative images of plaque assays performed using JEV viral samples from cells treated with NaFASO JINR1-S (control), NaFASO JINR1-1, NaFASO JINR1-2 respectively, showing variations in density corresponding to viral infectivity at different viral titter dilutions.
